# CRISPR-Cas systems in the plant pathogen *Xanthomonas* spp. and their impact on genome plasticity

**DOI:** 10.1101/731166

**Authors:** Paula Maria Moreira Martins, Andre da Silva Xavier, Marco Aurelio Takita, Poliane Alfemas-Zerbini, Alessandra Alves de Souza

**Affiliations:** Citrus Biotechnology Lab, Centro de Citricultura Sylvio Moreira, Instituto Agronômico de Campinas, Cordeirópolis-SP, Brazil; Departament of Microbiology, Instituto de Biotecnologia Aplicada à Agropecuária (BIOAGRO), Universidade Federal de Viçosa, Viçosa-MG, Brazil; Departament of Agronomy/NUDEMAFI, Universidade Federal do Espírito Santo, Brazil

**Keywords:** Phage, plasmids, *Xanthomonadaceae*, *Xylella*

## Abstract

*Xanthomonas* is one of the most important bacterial genera of plant pathogens causing economic losses in crop production worldwide. Despite its importance, many aspects of basic *Xanthomonas* biology remain unknown or understudied. Here, we present the first genus-wide analysis of CRISPR-Cas in *Xanthomonas* and describe specific aspects of its occurrence. Our results show that *Xanthomonas* genomes harbour subtype I-C and I-F CRISPR-Cas systems and that species belonging to distantly *Xanthomonas*-related genera in *Xanthomonadaceae* exhibit the same configuration of coexistence of the I-C and I-F CRISPR subtypes. Additionally, phylogenetic analysis using Cas proteins indicated that the CRISPR systems present in *Xanthomonas* spp. are the result of an ancient acquisition. Despite the close phylogeny of these systems, they present significant variation in both the number and targets of spacers. An interesting characteristic observed in this study was that the identified plasmid-targeting spacers were always driven toward plasmids found in other *Xanthomonas* strains, indicating that CRISPR-Cas systems could be very effective in coping with plasmidial infections. Since many effectors are plasmid encoded, CRISPR-Cas might be driving specific characteristics of plant-pathogen interactions.

## Introduction

Phytopathogenic bacteria are a global threat to crop production worldwide. *Xanthomonas* spp. is one of the most important genera of phytopathogens since these species can infect at least 120 monocotyledonous and 260 dicotyledonous species of economic importance (1,2). These pathogens are able to live both inside and outside of plant hosts. Regardless of their lifestyle, bacteria are constantly exposed to many different threats, such as the constant pressure in the form of exogenous DNA invasions from both viruses and invading plasmids from other bacteria (3,4). Many basic aspects of how these phytopathogens react and protect themselves from such threats remain understudied.

Bacteriophages (or simply “phages”) are one of the most abundant entities across the biosphere and one of the most potent pathogens of bacteria (5). Many aspects of both bacterial and phage genomes have been shaped by this never-ending war, in which both groups have had to develop defence and attack systems (6,7). In addition to virus attack, plasmid invasions can also be deleterious to bacteria. The most urgent topic concerning the negative effects of plasmidial invasions can be linked to the so-called “metabolic burden” (4,8), consisting of physiological disturbance due to the presence of exotic genetic material and its associated metabolism that drains important energetic resources of the host cell, negatively impacting its fitness.

For every horizontal genetic transfer that takes place in a prokaryotic cell, specific intra-cellular protection systems may come into play. Despite the fact that genomic rearrangements can lead to positive outcomes, there must be a balance between stability and tolerance of these events (3). Many biological systems have evolved to protect the integrity of the genetic information of prokaryotes. One of the first types of system ever discovered that eradicates exogenous DNA infections at their onset was restriction-modification systems, which recognize self-DNA by its methylation pattern and enzymatically destroy the invader DNA, thereby “restricting” its occurrence (9). Other mechanisms include the extreme abortive infection system, which kills the infected cell, preventing the phage from spreading throughout the bacterial population (10). Curiously, systems that were designed to aid in the maintenance of infective DNAs within cells have been co-opted for other functions. That is the case for the toxin-antitoxin operons (TA), which were originally described as a postsegregational killing system present in plasmids; infected cells that lose these invasive molecules will die, which increases plasmid prevalence among a given bacterial population (11). Few TA systems, such as the *mazEF* (12), *hok/soc* (13) and especially *toxIN* (14,15) systems, have been reported to exclude phages, mainly through the induction of premature cell death after phage invasion.

In the last decade, another bacterial defence system that has been in the spotlight is the CRISPR-Cas system (16). There are three types of CRISPR-Cas systems (I, II and III), each with many subtypes (17). These systems are basically characterized by the genomic presence of a module of repetitive DNA interpolated by “spacer” sequences consisting of previous invasive DNAs. During the occurrence of another invasion, these spacers are used to positively identify exogenous DNA and oppose the infective molecules (18). With the recent discovery of CRISPR-Cas as a defence mechanism in bacteria, its presence and abundance have been the focus of studies in the genomes of many prokaryotes, especially those of human-associated genera (19–22). However, in-depth analyses are lacking for phytopathogens, even in economically important genera such as the closely related taxa *Xanthomonas* and *Xylella fastidiosa* (23,24). In this work, we performed a genome-wide investigation of CRISPR-Cas in both of these phytopathogens, which cause diseases in different plant species, and showed that these systems may be a driving force for genetic diversity, impacting pathogenicity and host-range distribution.

## Materials and Methods

### Genome analysis

An in-depth analysis of both prophage and CRISPR arrays (and the identification of putative protospacer targets when CRISPR was present) was carried out in 10 *Xanthomonas* genomes that we previously selected (25). The complete list is shown in Table 1. The *Xylella fastidiosa* strains analysed for CRISPR arrays are also shown in Table 1, and subspecies were selected as phylogenetically representative members of these species (26). We expanded the number of genomes analysed only for the *cas* operon search to strengthen our conclusions about what subtypes of CRISPR-Cas systems are present in the *Xanthomonas* and *Xylella* genera. Therefore, the total numbers of genomes in this analysis were as follows: 121 *Xanthomonas* strains spanning 27 different species/pathovars (Supplemental File S1), 20 *Xylella* strains of four subspecies (Supplemental File 2S), and 7 other *Xanthomonadaceae* isolates (Supplemental File S3).

**Table 1:** Selection of genomes used for CRISPR array searches in both *Xanthomonas* spp. and *Xylella fastidiosa* ssp. genomes. (*) also used for prophage analysis.

| *Xylella fastidiosa* genomes | Accession number |
| --- | --- |
| *X. f.* subsp. *fastidiosa* Temecula | NC_004556.1 |
| *X. f.* subsp. *pauca* 9a5c | NC_002488.3 |
| *X. f.* subsp. *fastidiosa* M23 | NC_010577.1 |
| *X. f.* subsp. *multiplex* M12 | NC_010513.1 |
| *X. f.* subsp. *fastidiosa* GB514 | NC_017562.1 |
| *X. f.* subsp. *fastidiosa* MUL0034 | NZ_CP006740.1 |
| *X. f.* subsp. *sandyi* Ann-1 | AAAM04000275.1 |
| *Xanthomonas* genomes |  |
| * *X. axonopodis* pv. *citri* 306 | NC_003919.1 |
| * *X. axonopodis* Xac29-1 | NC_020800.1 |
| * *X. citri* subsp. *citri* Aw12879 | NC_020815.1 |
| * *X. campestris* pv. *vesicatoria* 85-10 | AM039952.1 |
| * *X. campestris* pv. *raphani* 756C | NC_017271.1 |
| * *X. campestris* pv. *campestris* ATCC33913 | NC_003902.1 |
| * *X. campestris* pv. *campestris* 8004 | NC_007086.1 |
| * *X. albilineans* GPE PC73 | NC_013722.1 |
| * *X. oryzae* pv. *oryzicola* BLS256 | NC_017267.2 |
| * *X. oryzae* pv. *oryzae* PXO99A | NC_010717.2 |

### CRISPR arrays and putative protospacer target identification

Based on the strain selection previously performed by Martins et al. (2016), who analysed the TA profiles of 10 *Xanthomonas* genomes that spanned the phylogenetic tree of this genus, we decided to use the same subset of *Xanthomonas* genomes to thoroughly analyse the origin of spacers. When a CRISPR array was identified, its putative protospacer targets were evaluated. For the *Xylella* genus, we selected 7 genomes spanning the four known subspecies (*fastidiosa*, *pauca*, *multiplex* and *sandyi*) regardless of their host range (Table 1). These genomes were submitted to CRISPR Finder (27) (http://crispr.i2bc.paris-saclay.fr/Server/), and the output of the CRISPR array when present was subsequently submitted to CRISPR Target (28) (http://bioanalysis.otago.ac.nz/CRISPRTarget/crispr_analysis.html) to identify possible matches to each of the spacers retrieved. The spacer content of every possible CRISPR array was analysed against the phage and plasmid databases provided by CRISPR Target, and to assess possible endogenous targets, we uploaded the bacterial genomes and performed the search again. In the case of a possible positive endogenous match, the sequences retrieved were further localized in the genome to identify the ORF. In these cases, the score threshold assumed for a positive ID was 5 mismatches (29). The full data for the targets identified are provided in Supplemental File S4. The results shown in Supplemental File S4 were classified into four colour-coded categories: unknown (pink), phage (green), plasmid (blue) and endogenous (yellow). To improve the analysis of the quantitative contribution of each of these targets them to the composition of the CRISPR array, the numeric values were submitted to the online CIRCOS Table viewer (30) (http://circos.ca/intro/tabular_visualization/).

### Prophage search

For the prophage search within the *Xanthomonas* genus, the same selection of 10 genomes used for the CRISPR array search was employed (Table 1) (25). The full chromosomal and plasmidial DNA sequences were submitted to the PHAST (31) and PHASTER (32) tools. Since our main objective was to assess viral infection entry (regardless of whether a full or incomplete prophage was involved), we used the PHAST output. The full PHAST output with the number of prophages found in each genome is shown in Supplemental File S5.

### *cas* operon search

Two different approaches were adopted to assess the presence of the *cas* operon in *Xanthomonas* and *Xylella*. The genomes already added to the CRISPI database (33) (http://crispi.genouest.org/) were analysed using this tool. However, the vast majority of the selected genomes are deposited as large contigs in the databases; in such cases, each CRISPR-Cas island was inspected and confirmed using CLC Genomics Workbench version 9.5.3 (QIAGEN) for the purpose of verifying the conservation of the *Cas* operon architecture.

CRISPR-Cas systems with acceptable CRISPR arrays and a Cas operon in the vicinity of these CRISPR units were considered valid(18). We considered CRISPR repeats embedded within ORFs to be false positives.

### Cas protein phylogeny

The Cas1 amino acid sequences of *Xanthomonadaceae* taxa from two CRISPR subtypes (I-C and I-F) were used for phylogenetic reconstruction along with Cas5d, Cas7 and Cas8c (CRISPR subtype I-C) downloaded from GenBank. The alignments were checked manually, and the evolutionary history was inferred using the maximum likelihood method based on the Jones-Taylor-Thornton (JTT) evolutionary model. Evolutionary analyses were conducted in MEGA7 (34). Additional phylogenetic trees using the Cas5d, Cas7 and Cas8c proteins (CRISPR subtype I-C) were generated to include the *Xylella taiwanensis* PLS229 taxon in this reconstruction since the operon in this isolate is eroded and does not contain the Cas1 protein, which is generally used in classical analyses.

## Results

### CRISPR repeat assessment

Some of the repeats reported by CRISPR Finder were considered false positives in our analyses. This was the case for one CRISPR locus from each the following 6 genomes*: X. citri* subsp. *citri* Aw12879, *X. citri* subsp. *citri* 306 and *X. axonopodis* Xac29-1, *X. campestris: X. campestris* pv*. campestris* ATCC 33913; *X. campestris* pv*. campestris* 8004 and *X. campestris* pv. *raphani* 756C. In all these cases, there was no associated *cas* operon in the vicinity of these repeats, and they were therefore considered false positives. These false-positive sequences and their repeats are shown in Supplemental File S6. No other CRISPR repeat region was dismissed, and they were all considered reliable. A summary of the final count of the number of CRISPR-Cas systems is shown in Supplemental File 8.

### The majority of *Xanthomonas* spp. present at least one *cas* operon

Among the twenty-seven different species/pathovars evaluated, 60% presented a putative functional CRISPR-Cas system (Supplemental File S7). For each given species/pathovar that showed at least one encoded *cas* operon, we observed that all of its strains presented the same type of *cas*. The only exception that we found was *X. oryzae* pv. *oryzicola*, in which one strain presented the subtype I-C system (str. YM15), while no CRISPR-Cas system was present in the other (str. BLS256) (Supplemental File S7). Among the CRISPR-Cas systems described to date (35), we only found Type I systems in *Xanthomonas*, of subtypes I-C and I-F (Supplemental File S6, Figure 1). The less prevalent subtype, I-F, was found exclusively in *X. fragariae*, *X. campestris* pv. *raphani* 756C, *X. albilineans* and *X. hyacinthi* (Figure 2A), while the most prevalent and widespread *Cas* operon found was I-C (Figure 2B), which was present in 15 of the 17 *Xanthomonas* species/pathovars with at least one *Cas* operon (Supplemental File S7). Curiously, *X. albilineans* and *X. hyacinthi* were the only species to present two *Cas* operons, each of which belonged to different subtypes: I-F and I-C. Interestingly, the same situation occurred in only two other *Xanthomonadaceae* species analysed as an outgroup (*Luteimonas huabeiensis* and *Dokdonella koreensis*) (Supplemental File S3, Figure 1); therefore, the presence of more than one Cas operon is a rare characteristic in this family. Curiously, none of the *Xylella fastidiosa* genomes analysed presented any CRISPR-Cas system, with the exception of *X. taiwanensis* PLS229, which presented a unique vestigial eroded subtype I-C-like *cas* operon, in which the genes encoding the Cas1, Cas2, Cas3 and Cas4 proteins were absent (Figure 2C).

**Figure 1.**
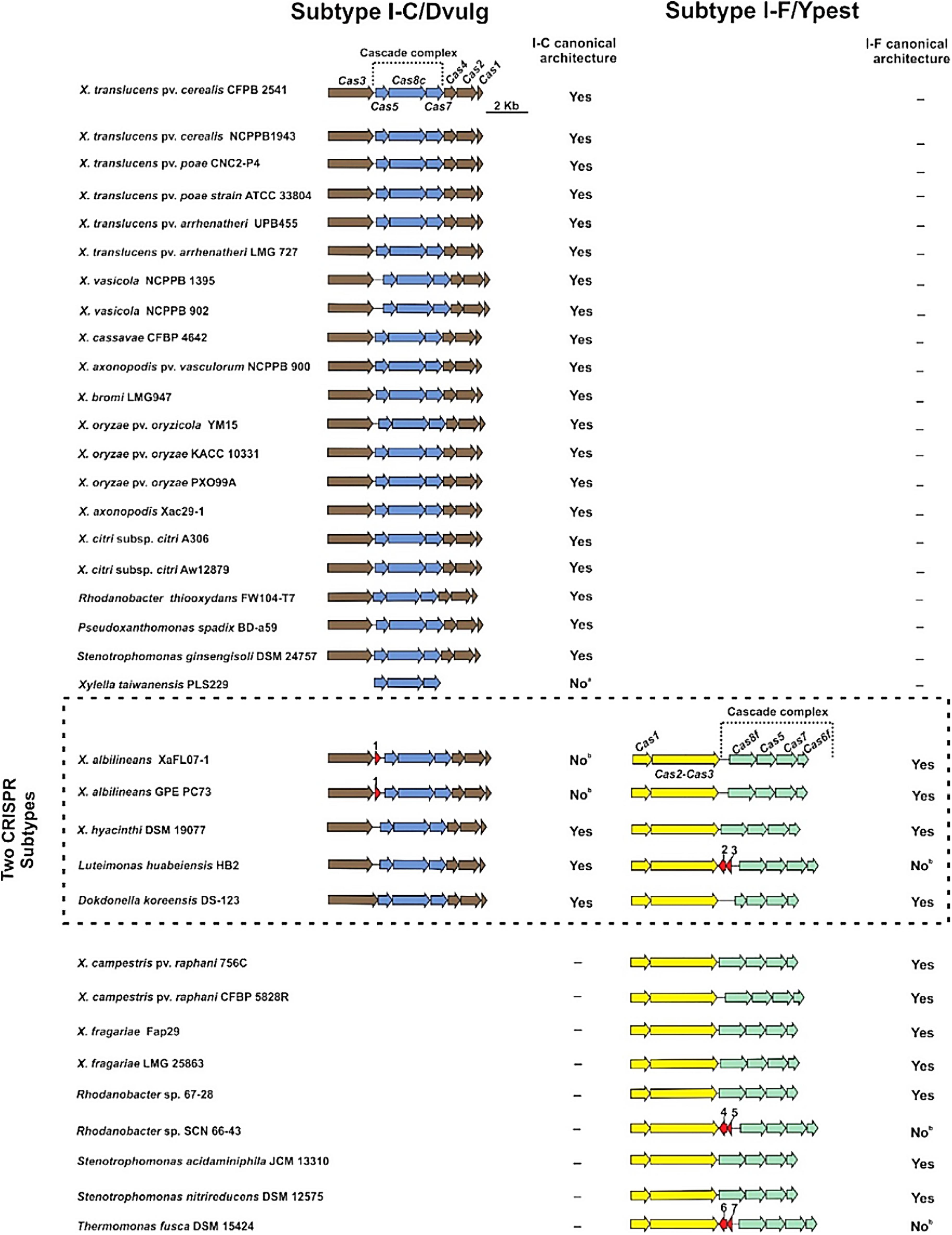
Overview of Cas operons found in *Xanthomonadaceae* species. *Xanthomonas* spp. predominantly carry Cas operons of subtype I-C, and Cas operons of subtype I-F can be found in some species at a lower frequency. Co-existence of the two CRISPR subtypes occurs in *X. albilineans* and *X. hyacinthi*. Taxa of other *Xanthomonas*-related genera also possess the two CRISPR subtypes and present a similar architecture, either occurring alone or co-existing in one isolate. Interestingly, some *Cas* operons do not possess the usual architecture and contain putative ORFs of completely unknown function in the stages of molecular execution by canonical CRISPR Type I. The region delimited by the dashed line indicates the species in which coexistence of the two CRISPR subtypes occurs. For the CRISPR I-C subtype, the adaptation and interference (cascade complex) modules are brown and blue, respectively, and for the CRISPR I-F subtypes, they are yellow and green, respectively. Unusual ORFs are represented as red arrows.

**Figure 2.**
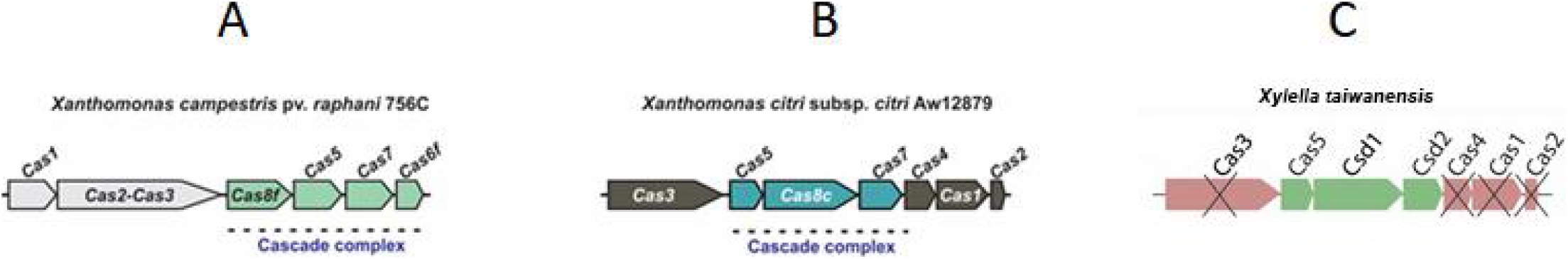
Cas operons found in *Xanthomonas* spp. A) *Xanthomonas campestris* pv. *raphani* is one of the strains to present only the I-F Cas operon subtype; B) *Xanthomonas citri* strains consistently presented a unique Cas operon of subtype I-C; C) Eroded Cas operon present in *Xylella taiwanensis*. The genes are depicted as arrows, with its putative gene names above. In green, the ORFs found in this genome, and with an “x”, the genes that are absent, but expected to be found in a subtype I-C CRISPR-Cas system.

### The Cas1 phylogeny shows ancestral acquisition of the Cas operon among *Xanthomonas* species

For both subtypes I-C and I-F, we observed a common phenomenon concerning the ancestrality of the Cas operon among *Xanthomonas* spp. strains (Figure 3A and 3B). Despite the wide range of horizontal gene transfer events in bacteria (36), it is reasonable to assume that Cas1 was acquired in a unique event of acquisition due to the high identity of this protein between different pathotypes of the same species. For instance, Figure 3A shows that *X. citri* 306 and *X. citri* Aw12879 exhibit identical Cas1 proteins despite their known differential host specificity (37). The same phenomenon was found in other species/pathovars showing different host preferences but presenting Cas1 clustering with high identity. The less prevalent subtype I-F showed exactly the same Cas1 profile, clustering the *Xanthomonas* strains of the same species/pathovar together (Figure 3B). Since no Cas operon or CRISPR was present in the *Xylella fastidiosa* genomes with the exception of *X. taiwanensis* PLS229, the phylogenetic reconstruction of the *Xanthomonadaceae* incorporating *X. taiwanensis* PLS229 was based on the sequences of the other proteins that are still present in this strain (Cas5d, Cas7/Csd1, Cas8c/Csd2) in an attempt to compare the phylogenetic signal of these unusual markers (Figure 4). The same taxon clusters detected in the phylogeny using Cas1 were observed in the trees generated using the Cas5d, Cas7/Csd1, and Cas8c/Csd2 proteins, in which the *Xanthomonas* strains of the same species/pathovars were clustered together, reinforcing the ancestral acquisition of the Cas operon among *Xanthomonas* species. For the *Xanthomonadaceae* analysed here, the phylogenetic signal of Cas1 (Figure 3) can be compared with those of other Cas proteins of subtypes I-C (Figure 4).

**Figure 3.**
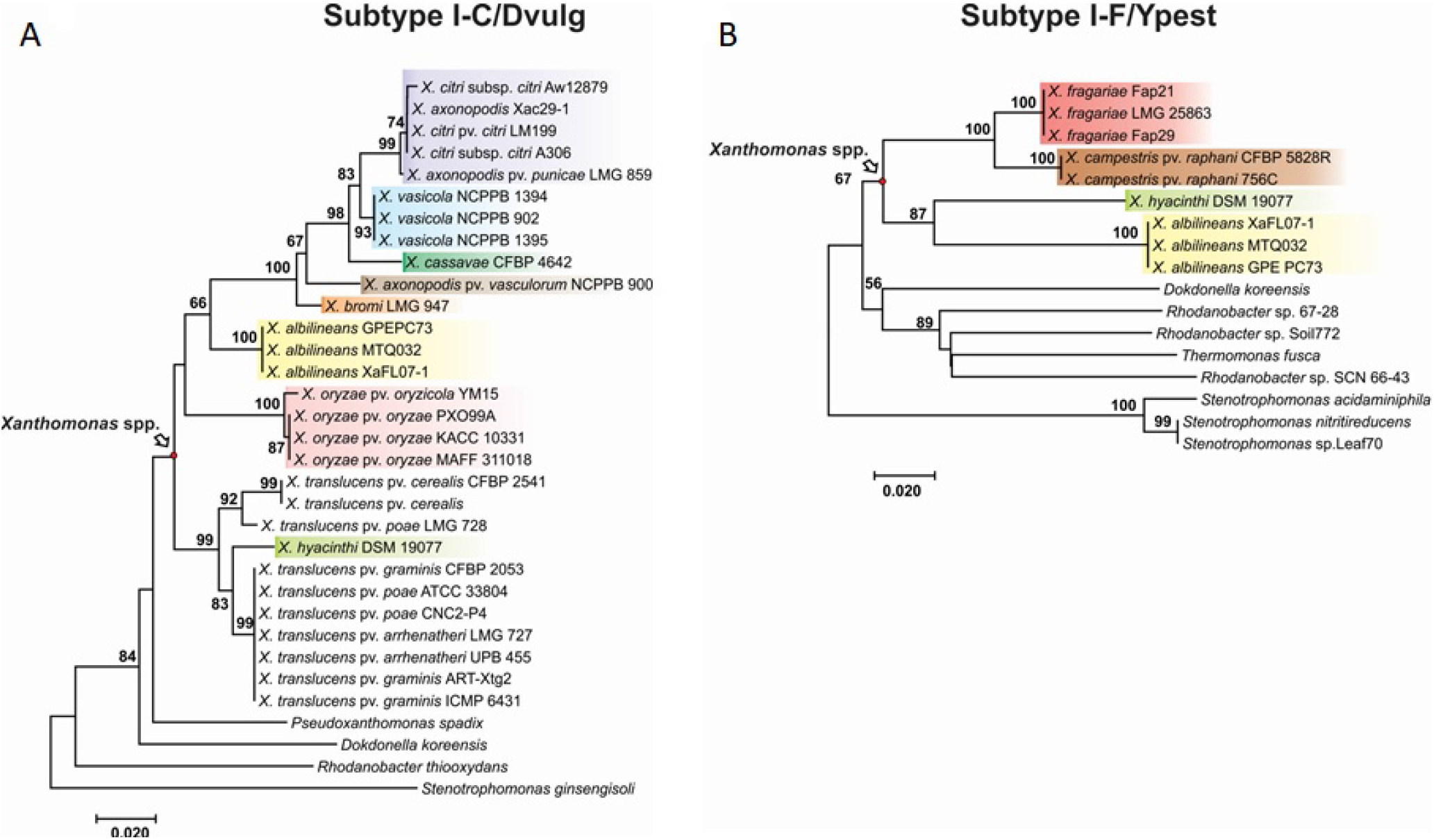
Cas1 phylogeny of subtypes I-F and I-C of CRISPR-Cas systems in *Xanthomonas* spp. and other *Xanthomonadaceae* species as outgroups. Highlighted coloured rectangles denote the clusters formed by the same *Xanthomonas* A) Subtype I-C, showing that *X. citri* strains cluster together despite having different hosts; B) subtype I-F, where the same phenomenon of species clustering despite different host preferences was observed. Bootstrap values (≥50%) are shown beside each node.

**Figure 4.**
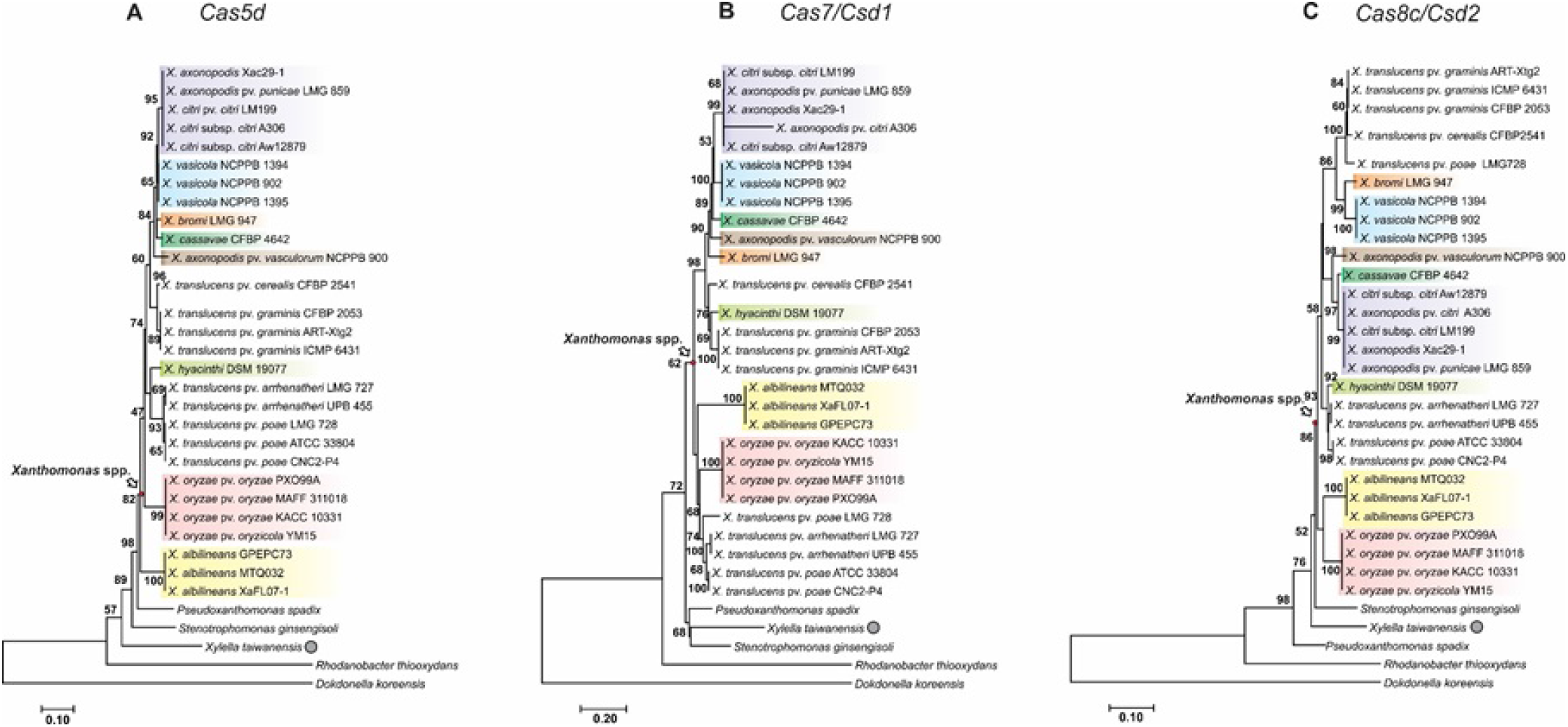
Cas5d, Cas7/Csd1 and Cas8c/Csd2 phylogeny of subtype I-C of CRISPR-Cas systems in *Xanthomonas* spp. and other *Xanthomonadaceae* species as outgroups. Highlighted coloured rectangles denote the clusters formed by the same *Xanthomonas* species/pathovars. The only species of *Xylella* that contains an eroded subtype I-C-like CRISPR-Cas system, *X. taiwanensis*, is highlighted in the tree with a grey circle. Bootstrap values (≥50%) are shown beside each node.

### The analysis of spacers shows a wide variety of targets

Each CRISPR locus of the strains was thoroughly analysed to assess the targets of each spacer. Although 40% of the strains showed no CRISPR-Cas systems whatsoever, those that harboured at least one such system showed variation in the number of spacers and their targets. The greatest number of these systems was found in *X. campestris* pv*. raphani* 756C (99 spacers), followed by *X. oryzae* pv. oryzae PXO99A (75 spacers). Additionally, although most of the targets were unknown, those that were identified showed matches with phage, plasmid and endogenous genome sequences (Figure 5).

**Figure 5:**
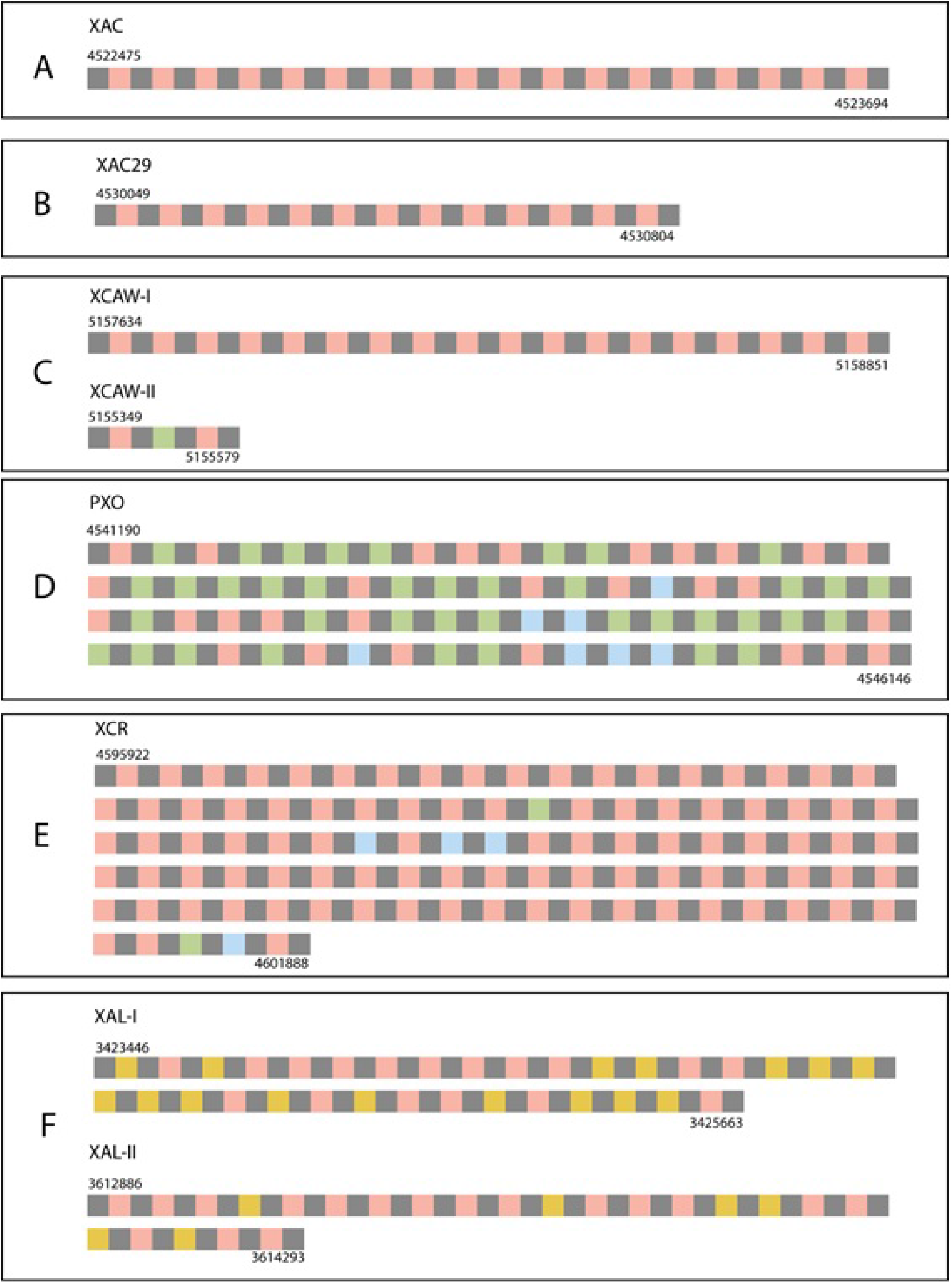
Schematic representation of CRISPR repeats and their spacers for each CRISPR locus. Grey squares represent repeats; yellow, pink, green and blue squares represent spacer targets of endogenous, unknown, phage and plasmid sequences, respectively. Numbers I and II are used to identify each CRISPR locus when more than one is found in the same genome. The genomic coordinates are indicated with the numbers above and under the squares. A) XAC (*Xanthomonas* citri subsp. citri 306); B) XAC29 (*Xanthomonas axonopodis* Xac29-1); C) XCAW (*Xanthomonas citri* subsp. *citri* Aw12869); D) PXO (*Xanthomonas oryzae* pv. *oryzae* PXO99A); E) XCR (*Xanthomonas campestris* pv. *raphani* 756C); F) XAL (*Xanthomonas albilineans* GPE PC73).

Our study showed that *X. oryzae* pv. *oryzae* PXO99A presented the greatest number of spacers targeting phages (frequently OP2, OP1 and Xop144) (Figure 5, green squares). In addition, spacer sequences targeting plasmids (Figure 5, blue squares) were found in *X. oryzae* pv. oryzae PXO99A and *X. campestris* pv*. raphani* 756C. Both *X. citri* subsp. *citri* 306 and *X. axonopodis* Xac29-1 presented one CRISPR-Cas system. However, the target could not be identified for any of the spacers encoded by their genomes. Likewise, *X. citri* subsp. *citri* Aw12869 presented one CRISPR-Cas system whose targets were not identified; however, a second CRISPR array was also detected in this strain. Despite the lack of an association with a *cas* operon in its vicinity, one of the spacer targets was positively identified in a phage sequence (Supplemental File S4). We therefore considered this second CRISPR array to be a putatively functional CRISPR-Cas system that may operate with the Cas proteins produced *in trans* by the first CRISPR-Cas system. Curiously, spacers targeting endogenous genome sequences were found at the two CRISPR loci only in *Xanthomonas albilineans* GPE PC73 (Figure 5, yellow squares). The percent contribution of each type of target in each CRISPR array is presented in Figure 6. The vast majority of unidentified matches and how the abundance of each category of spacers varies among the *Xanthomonas* strains are notable.

**Figure 6.**
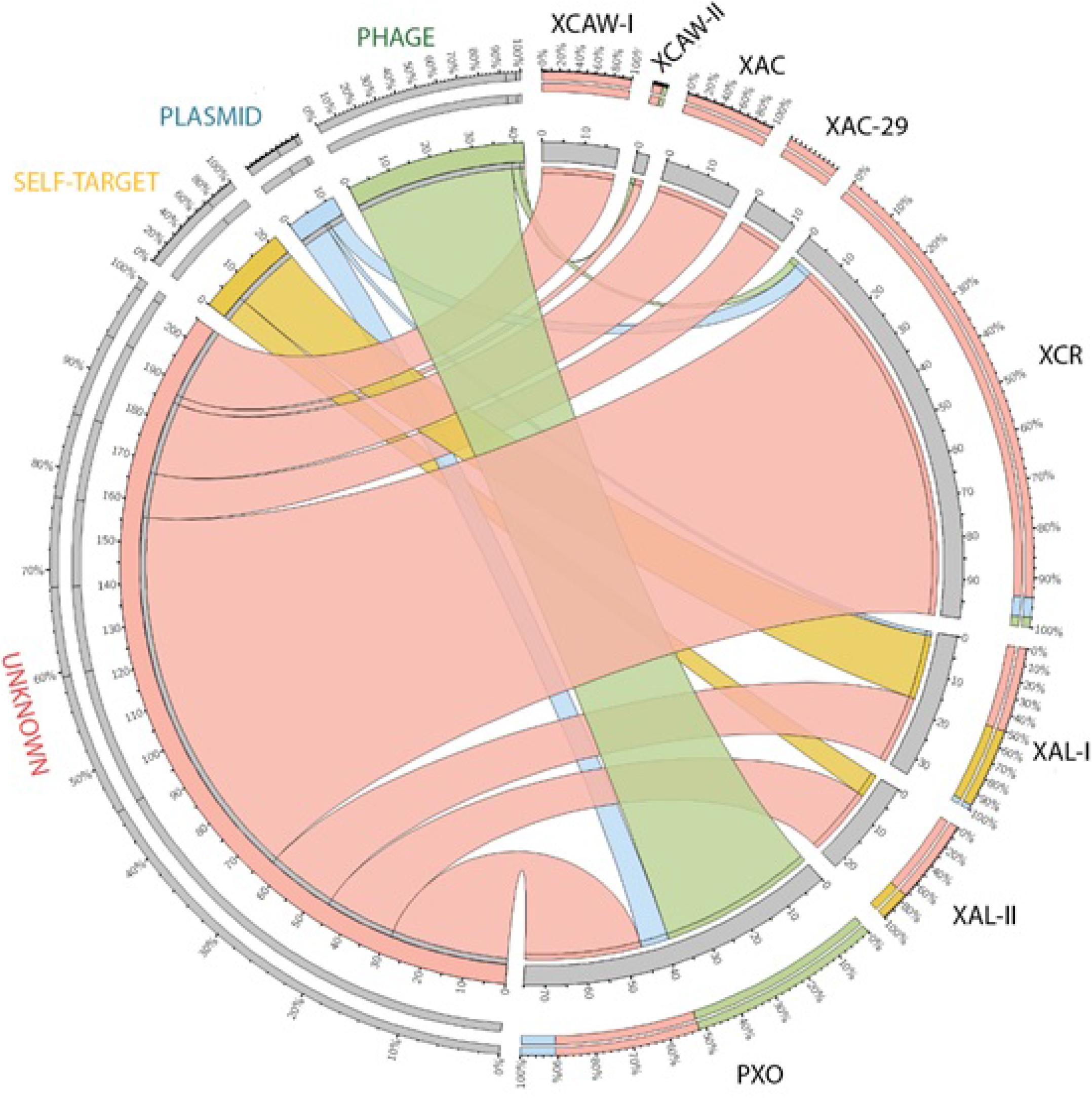
Percent contribution of each spacer target for each *Xanthomonas* spp. Ribbon colours represent the following categories pink: unknown; yellow: self-targets; green: phage-related; blue: plasmids. Numbers I and II are used to identify each CRISPR locus when more than one is found in the same genome. XAC (*Xanthomonas citri* subsp. *citri* 306); XAC29 (*Xanthomonas axonopodis* Xac29-1); XCAW (*Xanthomonas citri* subsp. *citri* Aw12869); PXO (Xanthomonas oryzae pv. oryzae PXO99A); XCR (*Xanthomonas campestris* pv. *raphani* 756C); XAL (*Xanthomonas albilineans* GPE PC73). Despite being less frequent, the plasmid targets that we identified were all from *Xanthomonas* plasmids.

### Spacers targeting *Xanthomonas* plasmids and prophage analyses

In addition to phages, plasmids targeted by the CRISPR-Cas systems were found in some *Xanthomonas* CRISPRs, and it was noteworthy that all of them presented the best matches to common *Xanthomonas* plasmids with high identity. Plasmids from *X. axonopodis* and *X. fuscans* subsp. *fuscans* are targets of the CRISPR-Cas systems of *X. campestris* pv*. raphani* (Figure 7 A, B and C). The other three spacers found in *X. oryzae* exhibited matches with 100% identity to plasmid targets of *X. citri* subsp. *citri* (Figure 7 D, E and F). We observed that *Xanthomonas* strains with more spacers presented fewer plasmids (Table 4). On the other hand, we noted that the *Xanthomonas* strains with many spacers targeting phages were those with more prophages integrated in their genome (Table 2).

**Figure 7.**
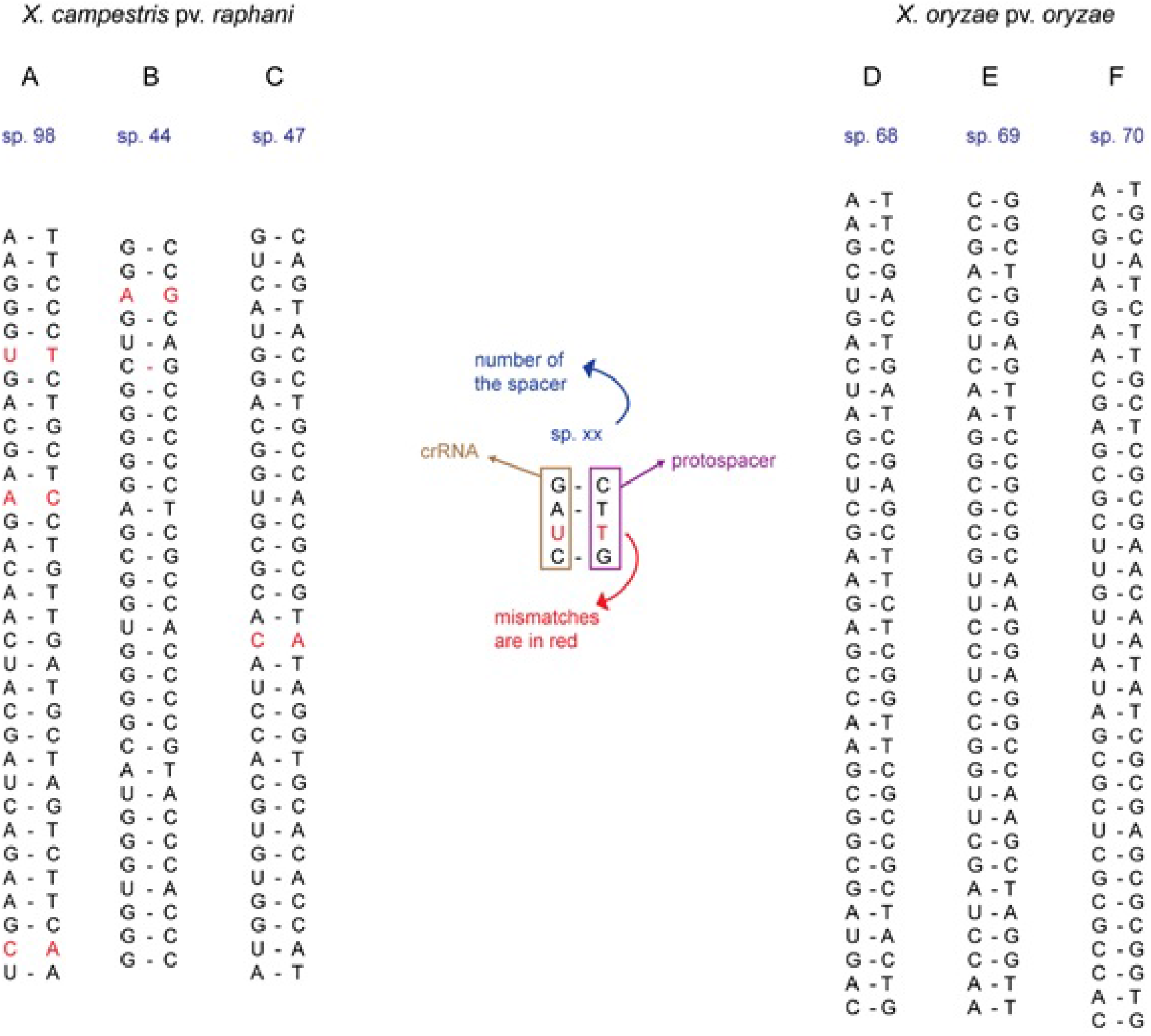
Selected examples of representative alignments between the putative transcribed crRNA and the protospacer. Alignments A, B and C are from spacers found in *X. campestris* pv. *raphani* 756C, and alignments D, E and F are from spacers found in *X. oryzae* pv. *oryzae* PXO99A. The protospacer identities are as follows: A) *X. axonopodis* Xac29-1 plasmid pXAC47; B) *X. fuscans* subsp. *fuscans* 4834-plasmid pla; C) *X. fuscans* subsp. *fuscans* 4834-plasmid pla; D) *X. citri* subsp. *citri* MN12 plasmid pXAC64; E) *X. citri* subsp. citri A306 plasmid pXAC64; F) *X. citri* subsp. citri NT17 plasmid pXAC64

**Table 2.**
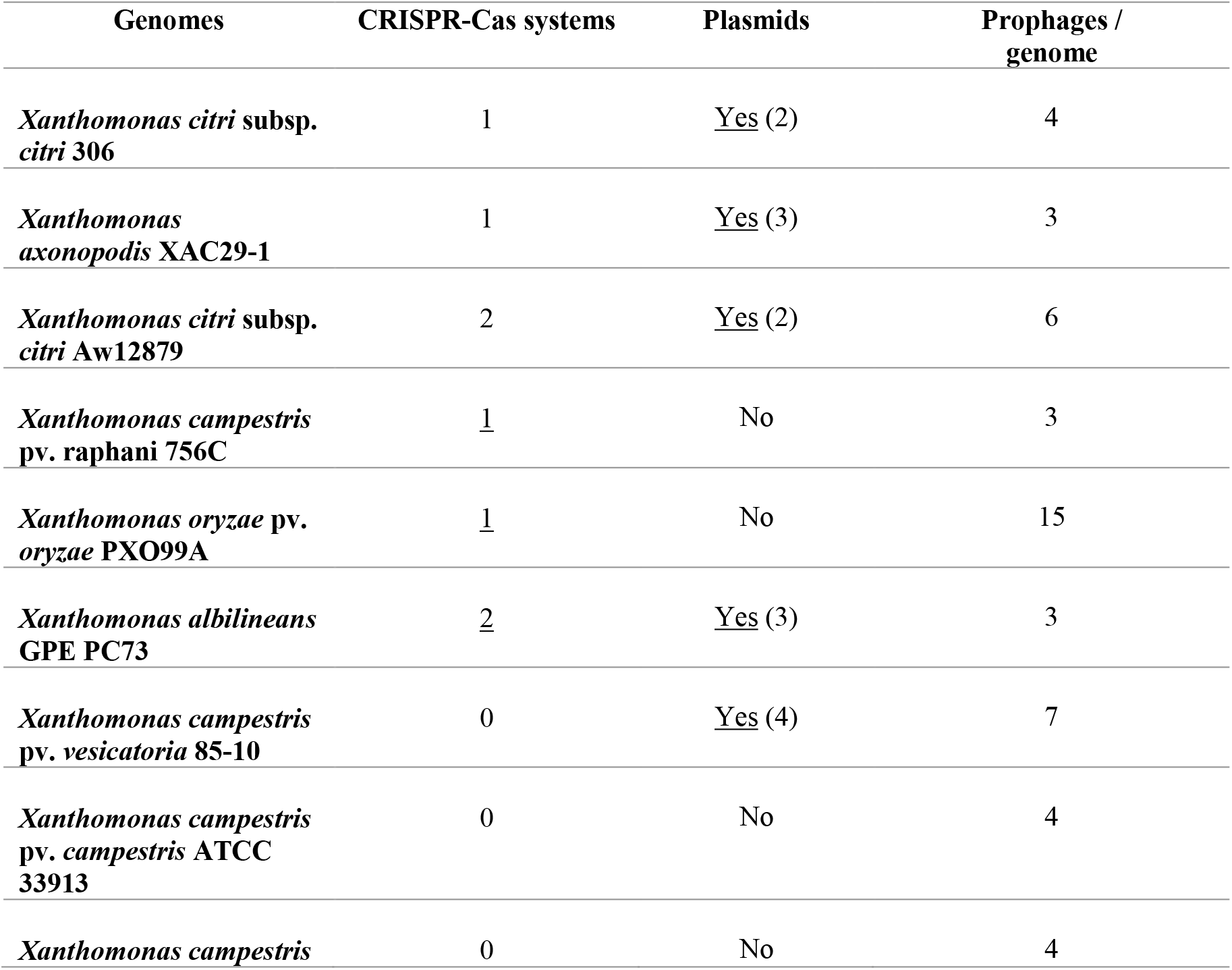
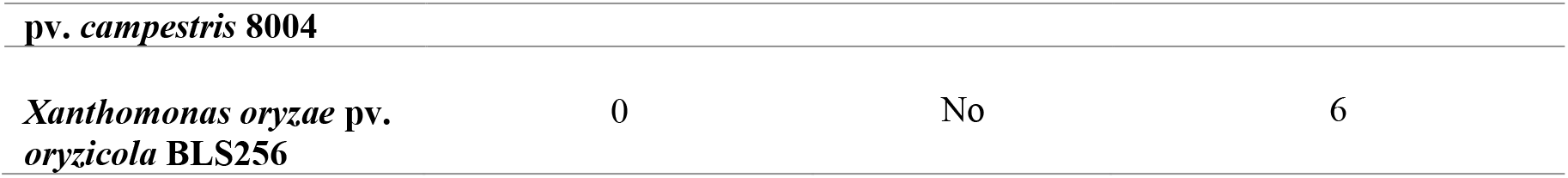
Presence of CRISPR-Cas versus mobile genetic elements. For the complete data, please see Supplemental Files S5 and S8

## Discussion

Recently, the CRISPR-Cas system was identified as a defence mechanism in bacteria; however, very little is known about its occurrence, abundance and targets in phytopathogens. Therefore, in this study, we performed a broad genome analysis of CRISPR-Cas in two closely related economically important genera, *Xanthomonas* and *Xylella*. Interestingly, no CRISPR-Cas system was found in *X. fastidiosa*. However, an eroded *Cas* operon was found in *X. taiwanensis*, a distant relative from Taiwan (26), raising the possibility that CRISPR-Cas systems may have been acquired but did not remain over time. It has been reported that entire bacterial lineages may lack CRISPR-Cas systems, as is the case for the *Chlamydiae* phylum among others, which is probably due to the potential deleterious autoimmunity risk that carrying a CRISPR-Cas system may pose (38). In addition, this absence may be a characteristic that is restricted to *Xylella* spp. and is not widespread among the *Xanthomonadaceae* since we also analysed 4 other genera within this family (*Thermomonas*, *Stenotrophomonas*, *Pseudoxanthomonas* and *Luteimonas*), and all of them showed at least one *Cas* operon. It is important to consider that phage-related regions of *Xylella fastidiosa* genomes can account for as much as 15% of the genome (39), which might be a direct result of the absence of CRISPR-Cas systems.

In contrast to the absence of CRISPR-Cas systems in *Xylella*, 60% of *Xanthomonas* spp. showed at least one *Cas* operon. Among the three types of CRISPR-Cas systems and their multiple subtypes (28), only Type I was identified in *Xanthomonas* (subtypes I-C and I-F). Usually, only one subtype was present per genome, except in *X. albilineans* and *X. hyacinthi*, which presented one copy of each I-C and I-F. We observed that the subtypes present in *Xanthomonas* (I-C, I-F or both together) were consistent among the strains of a particular species, which is in accordance with the observation that strains belonging to the same species usually harbour the same CRISPR-Cas system (40). We also highlight that species belonging to distantly *Xanthomonas*-related genera in *Xanthomonadaceae* presented the same configuration of coexistence of the same I-C and I-F CRISPR subtypes. In addition, our phylogenetic analysis indicated that the CRISPR systems present in *Xanthomonas* spp. are the result of an ancient acquisition.

Despite the similarities of the CRISPR-Cas subtypes in *Xanthomonas* spp. genomes, they presented significant variation in both the number and targets of spacers. The greatest number of spacers targeting sequences was observed in *X. oryzae* pv. *oryzae* PXO99A, which was in agreement with other studies that have emphasized the abundance of spacers in the CRISPR arrays of *X. oryzae* pv. *oryzae* (41,42). Regarding targets, self-targeting endogenous spacers were found only in *X. albilineans*, despite their presence in many bacterial genomes (43). The presence of self nucleic acids in CRISPR arrays indicates a form of autoimmunity that could explain the abundance of degraded CRISPR systems across prokaryotes (37). In addition, endogenous CRISPR spacers have been associated with regulatory mechanisms for repressing phage replication (44)). However, an important characteristic observed in this study was that the identified plasmid-targeting spacers were always driven toward plasmids found in other *Xanthomonas* strains, with *X. oryzae* pv. *oryzae* harbouring many of these spacers and being devoid of any extrachromosomal DNA. The same was true for *X. campestris* pv. *raphani*, which raises the possibility that CRISPR-Cas systems could be very effective in coping with plasmidial infections. Indeed *X. campestris* pv. *vesicatoria* and *X. fastidiosa* present more than one plasmid and no functional CRISPR-Cas system. On the other hand, the strain harbouring the greatest number of spacers targeting phages, *X. oryzae* pv. oryzae PXO99A, exhibited the greatest number of prophages in the genome, which may be a result of a very challenging environment concerning phage diversity but may also indicate that this system may not be functioning at the same rate at which viruses evolve to evade it (45). Therefore, CRISPR-Cas systems in *Xanthomonas* seem to be very effective in controlling plasmid infections, but they do not show the same success regarding phages. Since many effectors are plasmid encoded, CRISPR-Cas might be driving the specific characteristics of plant-pathogen interactions.

This is the first genus-wide analysis of CRISPR-Cas systems in *Xanthomonas*, and we conclude that the presence or absence of functional CRISPR-Cas systems may be an important driving force of genetic diversity in this genus, either allowing the entry and maintenance of DNAs in the cell or not, which may impose important gene flow restrictions in the course of evolution, consequently impacting the pathogenicity and host-range distribution of *Xanthomonas* spp.

## Authors statements

The authors declare the absence of any potential conflict of interest.

## Authors contribution

PM conceived and wrote the manuscript, and PM and AX executed the bioinformatics analysis. MAT, PAZ and AADS discussed the results and critically reviewed the manuscript.

## Acknowledgements

This work was supported by research grants from the Fundação de Amparo à Pesquisa do Estado de São Paulo (FAPESP - 2013/10957-0) and INCT-Citrus (CNPq 465440/2014-2 and FAPESP 2014/50880-0). PM is a FAPESP post-doctoral fellow (2016/01273-9).

